# Ring canals in the larval adipose of *Drosophila* buffer stress response

**DOI:** 10.1101/2025.06.11.658881

**Authors:** Shyama Nandakumar, Deepika Vasudevan

## Abstract

Cells in metabolically active tissues with high biosynthetic and secretory demands often use robust stress-responsive mechanisms to maintain endoplasmic reticulum (ER) homeostasis. Coordinating such robust stress response mechanisms requires intercellular communication and coordination. Such modalities of intercellular communication have been relatively understudied in the context of stress tolerance. Here, we use the *Drosophila melanogaster* third instar fat body to demonstrate that adipocytes communicate with each other through intercellular bridges called ring canals to buffer endoplasmic reticulum stress. The fat body supports the exponential growth from embryo to late larval stage over a short period of time through its energy storage and secretory functions, enduring a high basal level of ER stress in the process. We discovered that individual cells in the fat body are paired to one neighboring cell through ring canals. We further demonstrate that ring canals mediate rapid and highly specific intercellular cargo and organellar trafficking, and allow the transport of cytoplasmic, ER-bound and Golgi vesicular proteins. Disrupting fat body ring canals resulted in higher levels of ER stress response markers, aberrant cell size, as well as increased cell lethality in response to exogenous stress. We also find that animals with disrupted fat body ring canals display an overall delay in larval development, likely due to reduced secretion of larval serum proteins from the fat body. In sum, our work reveals a novel feature of intercellular communication in adipose tissue that serves to buffer ER stress across cells which is required for both homeostatic secretory function and maintaining tissue viability under exogenous stress.

## Introduction

Metabolically active tissues such as adipocytes and hepatocytes have high biosynthetic and secretory burdens. Due to their secretory demands, they also rely heavily on pathways that maintain endoplasmic reticulum (ER) homeostasis, such as the unfolded protein response (UPR) for their function[Han and Kaufman, 2016; Ryoo, 2024]. However, chronic activation of stress response pathways can result in pathologies, metabolic dysfunction, and ultimately, cell death. Thus, in order for a tissue to function properly under basal stress levels without invoking pathological stress programs, proper coordination and intercellular communication within a tissue is required. Intercellular communication within a tissue can be achieved through a number of avenues including short-range (physical connections such as cell-cell junctions[Cavey and Lecuit, 2009]), and cytonemes[González-Méndez et al., 2019]) and long-range methods (such as secreted molecules and hormones[Nässel and Zandawala, 2020]). How and whether these intercellular communication modalities support tissue-wide stress tolerance is relatively understudied.

Intercellular bridges are formed at the end of a cell division cycle when, following an incomplete cytokinesis, the cytokinetic furrow is stabilized to form a stable connection between two daughter cells[Chambaud et al., 2024; Lu et al., 2017]. Such structures are well conserved across multicellular organisms, albeit known by different names across popular model organisms; septin rings in yeast[Kukhtevich et al., 2024], ring canals in *Drosophila*[Robinson and Cooley, 1996; Robinson et al., 1994], intercellular bridges in mammals and other vertebrates[Andreuccetti et al., 1999; Fawcett et al., 1959; Mullins and Biesele, 1973], and as plasmodesmata in plants[Cilia and Jackson, 2004; Zani and Edelman, 2010]. However, the structural components of plasmodesmata differ from intercellular bridges, septin rings, and ring canals found in metazoans as they traverse cell walls. Importantly, in each of these cases, intercellular bridges serve as channels of active, highly regulated transport and intercellular communication, and not merely diffusion.

The function of intercellular bridges in animals is best studied in the context of the germline [Airoldi et al., 2011; Kloc et al., 2004; Lee et al., 2013; Price et al., 2023; Roth and Lynch, 2009]. In *Drosophila*, ring canals have several known functions depending on the sex of the germline. In the female germline, they allow trafficking of maternally loaded mRNAs and proteins synthesized in the nurse cells into the developing oocytes[Bernard et al., 2018; Hsu et al., 2015; McLean and Cooley, 2014; Petrella et al., 2007; Roth and Lynch, 2009; Shaikh et al., 2024]. In the male germline, they function to synchronize meiotic division and development of male spermatids in the testes[Eikenes et al., 2015; Kaufman et al., 2020; Miyauchi et al., 2013; Montembault et al., 2010]. Intercellular bridges have been described to perform similar, conserved functions in the mammalian germline as well (reviewed in [Greenbaum et al., 2007, 2011]). While the roles of intercellular bridges have been well established and described in the germline, the role that these structures may play in somatic tissues is relatively unknown. Ring canals have been described in a few other somatic tissues such as the *Drosophila* wing disc in larvae[McLean and Cooley, 2014], main cells of the male accessory glands[Box et al., 2024] and in follicle cells of adult female ovary[Airoldi et al., 2011; McLean and Cooley, 2013], but their specific function in each of these tissues remains unstudied.

Further, the composition and size of the intercellular bridges is cell type dependent. For example, in *Drosophila* the largest ring canals connect nurse cells in the female germline, can be anywhere between 1-10µm in diameter depending on developmental stage, and these ring canals contain actin, among dozens of other structural proteins[Hudson and Cooley, 2002; Kelso et al., 2002; Warn et al., 1985]. In contrast, the male germline ring canals do not contain actin, and are up to 4µm in diameter.[Gerdes et al., 2020; Haglund et al., 2011; Hime et al., 1996] Notably *Drosophila* ring canals have been demonstrated to require several protein complexes to maintain their ‘openness’, including actin remodeling complexes, plasma membrane bound regulators of signaling pathways such as src kinases, and proteosome-associated proteins such as the E3 ubiquitin ligase, Kelch [Cooley, 1998; Guarnieri et al., 1998; Robinson and Cooley, 1996; Robinson et al., 1994] .

The *Drosophila* adipose tissue, also called the ‘fat body’, has several vital secretory, endocrine, innate immune and lipid storage functions[Tennessen and Thummel, 2011]. In addition to these functions, the larval fat body supports the exponential growth of the embryo to pupa over a short developmental period of 120h [Church and Robertson, 1966]. This tissue is derived from mesodermal progenitors which stop proliferating in embryonic stages [Hartenstein and Jan, 1992; Moore et al., 1998; Riechmann et al., 1997] to give rise to the postmitotic fat body. To support the exponential growth of the larvae through development, adipocytes themselves undergo multiple rounds of endocycling, or truncated cell cycles which involve alternating growth and DNA synthesis phases to grow in cell size, but not in number. [Colombani et al., 2003; Pierce et al., 2004; Smith and Orr-Weaver, 1991]. By the last larval instar stage, the fat body is organized as multiple lobes across the entire body plan of *Drosophila*, comprising approximately ∼2,000 cells each with an average ploidy of 255C [Nordman et al., 2011]. Across all these stages, but particularly in the third instar stage, the fat body secretes vast amounts of proteins and lipids into the hemolymph to support the growth of all the developing tissues, as well as in preparation for pupariation[Heier et al., 2021; Jowett et al., 1986; Powell et al., 1984; Valzania et al., 2024]. The secretory profile of the fat body includes a class of proteins called larval serum proteins (Lsps), which are taken up by various tissues, for use as energy reserves during metamorphosis during which animals neither move nor feed. [Roberts et al., 1991; Wolfe et al., 1977].

Tissues with high secretory activity, such as the *Drosophila* fat body, often display high levels of stress. Consistent with this, we and others have shown that the third instar larval fat body also displays elevated activation of ER stress markers [Huang et al., 2017; Kang et al., 2015, 2017; Sone et al., 2013]. There are three well-studied ER stress response pathways, each activated by an ER stress-sensitive protein - IRE1, PERK, and ATF6- which then signal through their downstream effectors- XBP1, ATF4, and cleaved ATF6 respectively[Walter and Ron, 2011]. Both XBP1 and ATF4 activity is present at high basal levels in the wandering third instar larval fat body, reflective of the ER stress burden in this tissue [Huang et al., 2017; Kang et al., 2015, 2017; Sone et al., 2013]. Whether and how the ER stress burden is coordinated across the tissue to cope with the constant secretory demand in this tissue has not been studied. Here, we show that individual adipocytes are paired to exactly one neighboring cell through a ring canal, rendering a tissue geometry where adipocytes form bi-cellular syncytia. We find that ring canals are present in both males and females, across all lobes of the fat body, and throughout all the larval stages of development. Using advanced imaging paradigms, we determine the dimensions of the ring canals as well as the specificity of cargo transport that they permit. Further we show that fat body specific knockdown of *kelch* alters the dimensions of ring canals and results in abrogated intercellular transport of cytoplasm. Additionally, compromising ring canal structure results in developmental delays and reduced secretion of larval serum proteins. Fat bodies with disrupted ring canals also show “unpairing” of ‘sister’ adipocytes resulting in large variation in stress reporter expression and cell sizes between pairs. Such unpairing also increases the susceptibility of the fat tissue to exogenous ER stress, indicating that, in this context, ring canals could serve to buffer cellular stress and promote cell survival under high secretory burden conditions. Together, our work is not only the first description of ring canals in adipocytes but also uncovers a novel function for ring canals in a somatic tissue with critical roles in development.

## Results

### Pairs of Drosophila larval adipocytes are connected via ring canals

When studying the stress tolerance of the fat body using a transgenic reporter that reads out the activity of PERK (4E-BP^intron^-DsRed, [Kang et al., 2017]) in the we observed a pattern where neighboring adipocytes appear to display similar levels of reporter gene expression (**Fig. S1**), suggesting that the cells may somehow be paired to one another. This pattern resembled the expression pattern of transgenes in other somatic tissues of *Drosophila* which have been shown to contain ring canals, such as the accessory glands [Box et al., 2024] in adult males and follicle cells in adult female ovaries [Airoldi et al., 2011; McLean and Cooley, 2013]. We thus sought to test if the neighboring adipocytes in the fat body were connected via ring canals using the pavarotti-GFP (pav-GFP) transgenic reporter [Airoldi et al., 2011]. Pavarotti is a kinesin family protein (with close homology to mammalian KIF23) which is known to localize to ring canals in many *Drosophila* tissues[Bassi et al., 2013; Carmena et al., 1998; Eikenes et al., 2013]. We found pav-GFP puncta throughout the wandering third instar fat body and present exclusively on the cell membrane (**Fig. 1**). We determined that individual adipocytes only have one ring canal puncta per cell, indicating that pairs of adipocyte nuclei are potentially connected through ring canals. We next examined whether such putative ring canals are present throughout the fat body.

**Figure 1:**
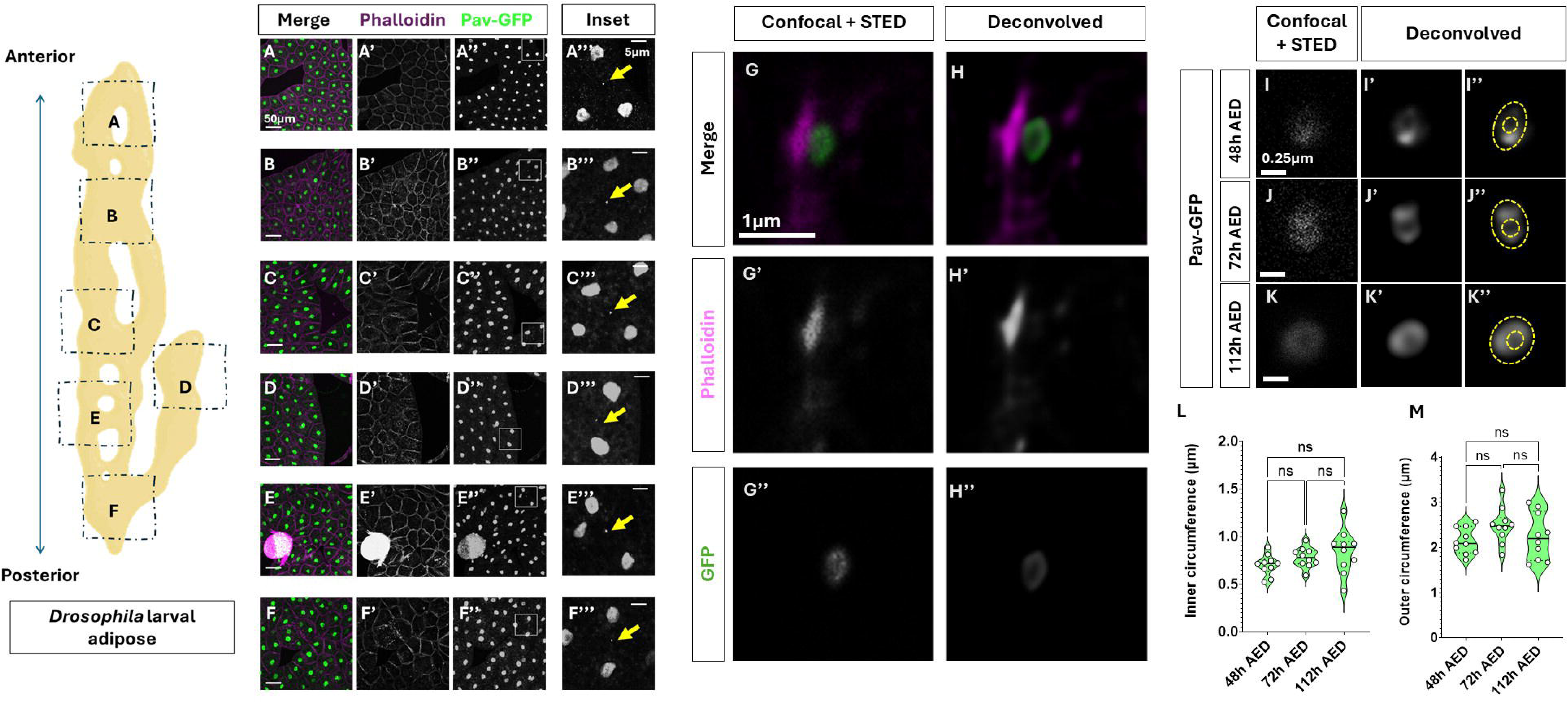
Super resolution microscopy reveals pav-GFP puncta found throughout the *Drosophila* larval fat body on adipocyte cell membranes and represent open ring canals. Schematic (left). Cartoon of the larval fat body (made using Biorender) displaying orientation of the *Drosophila* larval fat body with anterior-posterior orientation denoted. Dotted and labelled rectangles show the regions of the fat body that were imaged in panels A-F’’’. A-F. Confocal images of wandering third instar larval fat bodies expressing pav-GFP (green) and counterstained with Phalloidin (magenta) to mark cell membranes. Note that pav-GFP localizes to the nucleus, putative ring canals and midbody remnants. Inset regions marked by white rectangles in panels A’’-F’’ are magnified and shown in A’’’-F’’’. Yellow arrows in A’’’-F’’’ point to minute pav-GFP puncta between two adipocyte nuclei. G-H. Representative high magnification (93X Objective) confocal image of pav-GFP (green) puncta from Fig. 1A with STED super resolution (G-G’’). Cell membranes are counterstained with phalloidin (magenta). H-H’’’’ show deconvolved image from G-G”. I-K. STED super resolution and deconvolution confocal images of pav-GFP puncta from fat bodies at different developmental timepoints corresponding to 48h after egg deposition (AED) (I-I’’), 72h AED (J-J’’) and 112h AED (K’-K’’). Yellow dotted lines outline the inner and outer circumference of the ring canal in I”-K”. L-H. Measurements of the inner and outer circumference of ring canals from super-resolved images in C-E across larval development at indicated timepoints. n=10 ring canals from ten animals Scale bars: A-F, A’-F’, A’’-F’’ = 50µm; A’’’-F’’’ = 5µm; G-H’’ = 1µm; I-K = 0.25µm ns= not significant; ****=p<0.00001; ***=p<0.0001; *=p<0.01 here and in all other figures.

Using fluorescence confocal imaging, we determined that adipocytes in all lobes of the larval fat body are paired with exactly one neighbor via a pav-GFP puncta (**Fig. 1A-F**). While we observed ring canals in both male and female fat bodies, we chose to utilize females for the rest of this study.

### Ring canals are open structures and their size remains unchanged throughout larval development

With the goal of ultimately understanding the specificity of ring canal-mediated intercellular communication, we wanted to determine whether the pav-GFP puncta represented open ring-canals or simply midbody remnants. Midbody remnants are vestiges of cytokinesis, and sometimes remain intact in daughter cells after cell division has occurred [Price et al., 2022]. Due to their size, we were unable to resolve whether the GFP puncta represent open ring canals or midbody remnants even at the highest magnifications of traditional, high-resolution confocal microscopy (**Fig. 1G**). To understand the dimensions and structure of the puncta better, we turned to STED super resolution[Vicidomini et al., 2018] followed by Huygens deconvolution, which revealed less pixel dense center of pav-GFP puncta resembling open ring canal structure, on the cell membrane(**Fig. 1H**). Scaled measurements revealed that the open ring canals are approximately 1.8-3μm in outer circumference with a 0.5-1.2 μm inner circumference (**Fig. 1H, L,M**).

Since the larval fat body is post-mitotic and the number of adipocytes is determined at the end of embryogenesis [Pierce et al., 2004; Smith and Orr-Weaver, 1991][Butterworth et al., 1988; Edgar and Orr-Weaver, 2001], we surmised that ring canal formation must have occurred early in fat body development. Consistent with this, we found that ring canals are present in all in larval fat bodies from 48h AED (hours after egg deposition) to previously tested wandering third instar larva at 112h AED (**Fig. 1I-K**). We next measured the dimensions of ring canals across development from 48h AED to 112h AED which represents the majority of larval growth stages. Surprisingly, despite these stages coinciding with exponential endocycle-mediated growth of individual adipocytes [Guarner et al., 2017; Nordman et al., 2011; Pierce et al., 2004], we found that ring canals don’t change in size between 48h AED and 112hAED (**Fig. 1L-M**). These results strongly suggest that ring canal formation occurs as a result of an incomplete cytokinesis at the end of the last mitotic cell cycle in late embryogenesis [Pierce et al., 2004; Smith and Orr-Weaver, 1991].

#### Intercellular transport across ring canals shows cargo-bias

The maintenance of a relatively small ring canal size throughout larval stages of adipocyte cell and nuclear growth (**Fig. 1L-M**) suggests that ring canals allow for size-selective specific cargo transport. To test candidate cargos that may be transported across ring canals, we devised a fluorescence recovery after photobleaching (FRAP) experimental paradigm. We also developed a protocol for *ex vivo* live imaging of fat body from wandering third instar larva (see methods) expressing these fluorescently labeled organelles for FRAP experimentation. Using a fat body driver (*Dcg-GAL4*, [Asha et al., 2003]), we drove the expression of various compartment-bound fluorescent proteins in adipocytes. We first tested whether cytoplasmic proteins are able to pass through the ring canal using a cytoplasmic GFP. We photo-bleached cytoplasmic GFP in one adipocyte, and measured changes in fluorescence over time in the bleached cell, the neighboring cell paired through a ring canal, and an “unpaired” adjacent neighbor not connected to the cell through a ring canal (**Fig. 2A**). We observed that the bleached cell rapidly recovered GFP fluorescence in 30 seconds, with a concomitant decrease in fluorescence in the paired neighbor and no change in fluorescence in the unpaired neighboring cell (**Fig. 2A’, A”**). To ensure that such recovery in paired cells accounts only for ring canal mediated transport of cytoplasmic GFP and not de novo fluorescent protein biosynthesis, we photo-bleached both paired neighbors (**Fig. 2B**). Across similar time scales of imaging as **Fig. 2A**, we found that neither of the bleached pair showed any recovery in fluorescence during the course of imaging (**Fig. 2B’, B”**). These experiments demonstrate that cytoplasmic content is continually and rapidly exchanged between adipocytes connected by ring canals.

**Figure 2:**
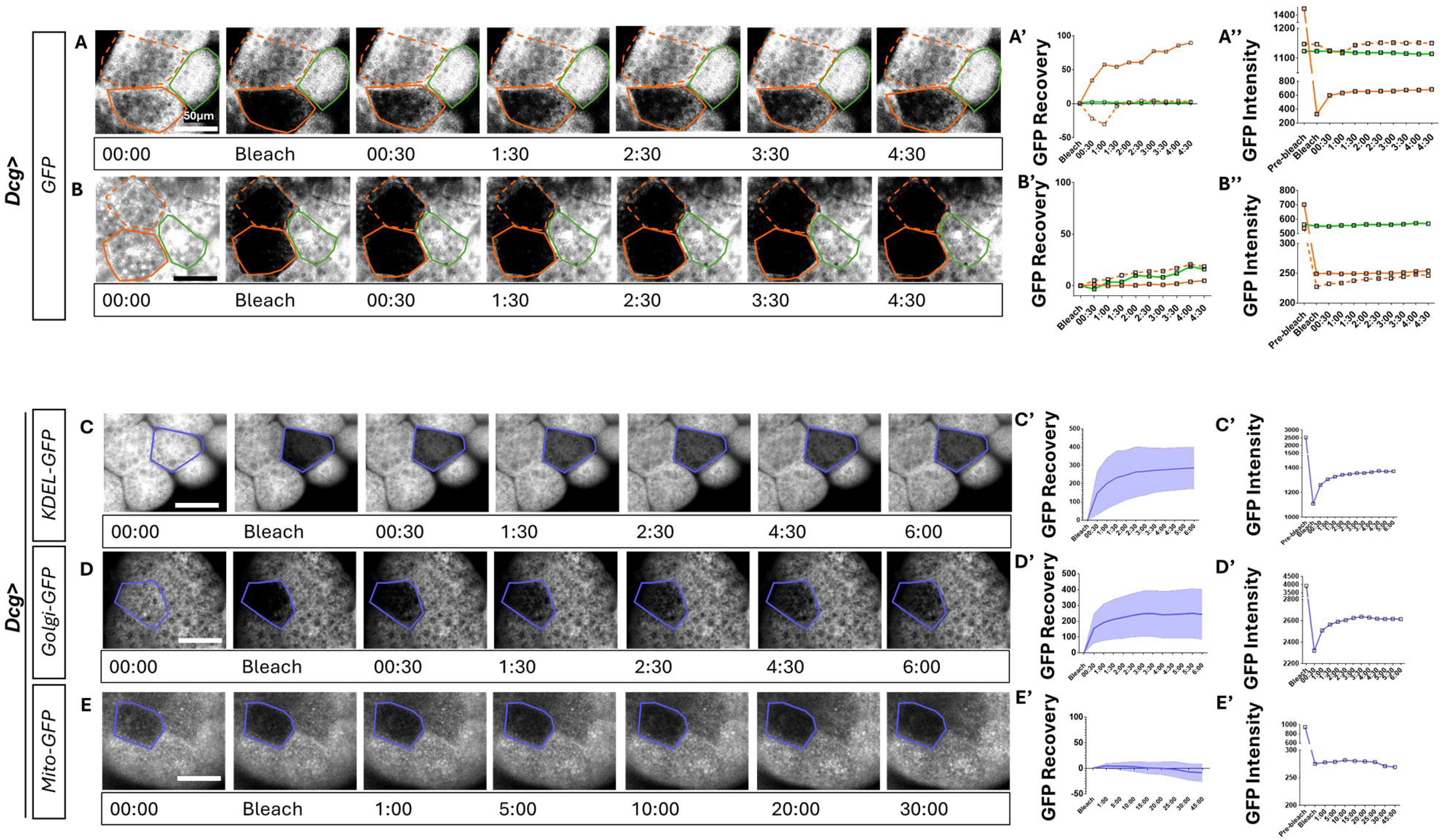
Cytoplasmic, ER and Golgi-bound fluorescent proteins are transported rapidly through rig canals, but not mitochondrial proteins. A-B. Fluorescent image montages showing time course of GFP signal over 4.5 minutes from *Dcg-GAL4>UAS-GFP* fat bodies subjected to photobleaching followed by recovery (FRAP) for several minutes. Panel A shows montage corresponding to one bleached cell and panel B shows experimental paradigm where both the adipocytes connected through ring canal are subjected to photobleaching (solid orange outlines). GFP fluorescence intensity is measured in the bleached cell (orange outline), paired neighboring cell (dashed orange outline) and a third neighboring cell (green outline) and plotted as recovery following photobleaching normalized to initial values in A’-B’ and as absolute values in A”-B”. C-E. FRAP analysis on ER-bound KDEL-GFP (C), Golgi bound-GFP (D) and mitochondria-GFP (E) as performed in A-B where GFP recovery is measured in the bleached cell (blue outline). Note that C-D were performed over a time course of six minutes and E was performed over 30 minutes. Quantifications in C’-E’ represent the mean change in GFP recovery post bleaching in the bleached cell. Standard deviation is plotted as the shaded error envelope. n=10 independent FRAP experiments from ten animals over 3 independent crosses. Quantification in C”-E” are absolute GFP intensity values from images in C-E. Scale bars for all panels = 50µm

Since the fat body is a highly metabolically active and secretory tissue, we next tested whether mitochondria, ER, or Golgi compartments maybe transported across ring canals. To do so, we used *Dcg-GAL4* to drive expression of either *UAS-KDEL-GFP* (marking the ER), *UAS-Golgi-GFP* (marking Golgi vesicles) or *UAS-mito-GFP* (marking mitochondria). Similar to the cytoplasmic GFP, we found that both KDEL-GFP and Golgi-GFP are transported rapidly (in under three minutes) across paired neighbors (**Fig. 2C-D**). In contrast, the mitochondrial GFP did not recover at even 30 minutes after photobleaching (**Fig. 2E**) suggesting that mitochondria likely do not get transported through ring canals in third instar larval adipocytes. These findings demonstrate that cargo transport across paired adipocytes is selective and permits transfer of secretory pathway components.

### Kelch is required for maintaining fat body ring canal integrity

Previous work from our lab and others has shown that the developing larval fat body endures high levels of intrinsic ER stress, which is a feature of its function [Kang et al., 2015, 2017; Sone et al., 2013]. Since we found intercellular exchange of secretory pathway components, we hypothesized that ring canals may play a role in regulating ER stress across paired adipocytes. To test this, we sought to disrupt ring canal structure and examine the consequences of such disruption. One of the best-described regulators of ring canal integrity in the female germline is the E3 Ubiquitin ligase, *Kelch* [Hudson et al., 2015, 2019; Kelso et al., 2002]. *Kelch* stabilizes ring canal structure by targeted substrate degradation to keep the ring canals open. To test whether adipocyte ring canal integrity is also regulated by *Kelch*, we performed fat body-specific knockdown of *Kelch* using the *Dcg-Gal4* driver (henceforth called *Dcg>Kelch^RNAi^*) and compared them to a control RNAi (*Dcg>LacZ^RNAi^*). Super-resolution STED, followed by deconvolution of ring canals from fat bodies from *Dcg>Kelch^RNAi^*wandering third instar larva revealed that loss of *Kelch* resulted in decreased inner circumference of ring canals but that the outer circumference of the ring canal was unaltered (**Fig. 3A-B, D-E**). We also observed similar reduction in inner circumference of the ring canals in *Kelch* mutant heterozygotes (*Kelch^DE1^/+,* [Hudson et al., 2019]) (**Fig. 3C-E**), corroborating our RNAi result [Hudson et al., 2019].

**Figure 3:**
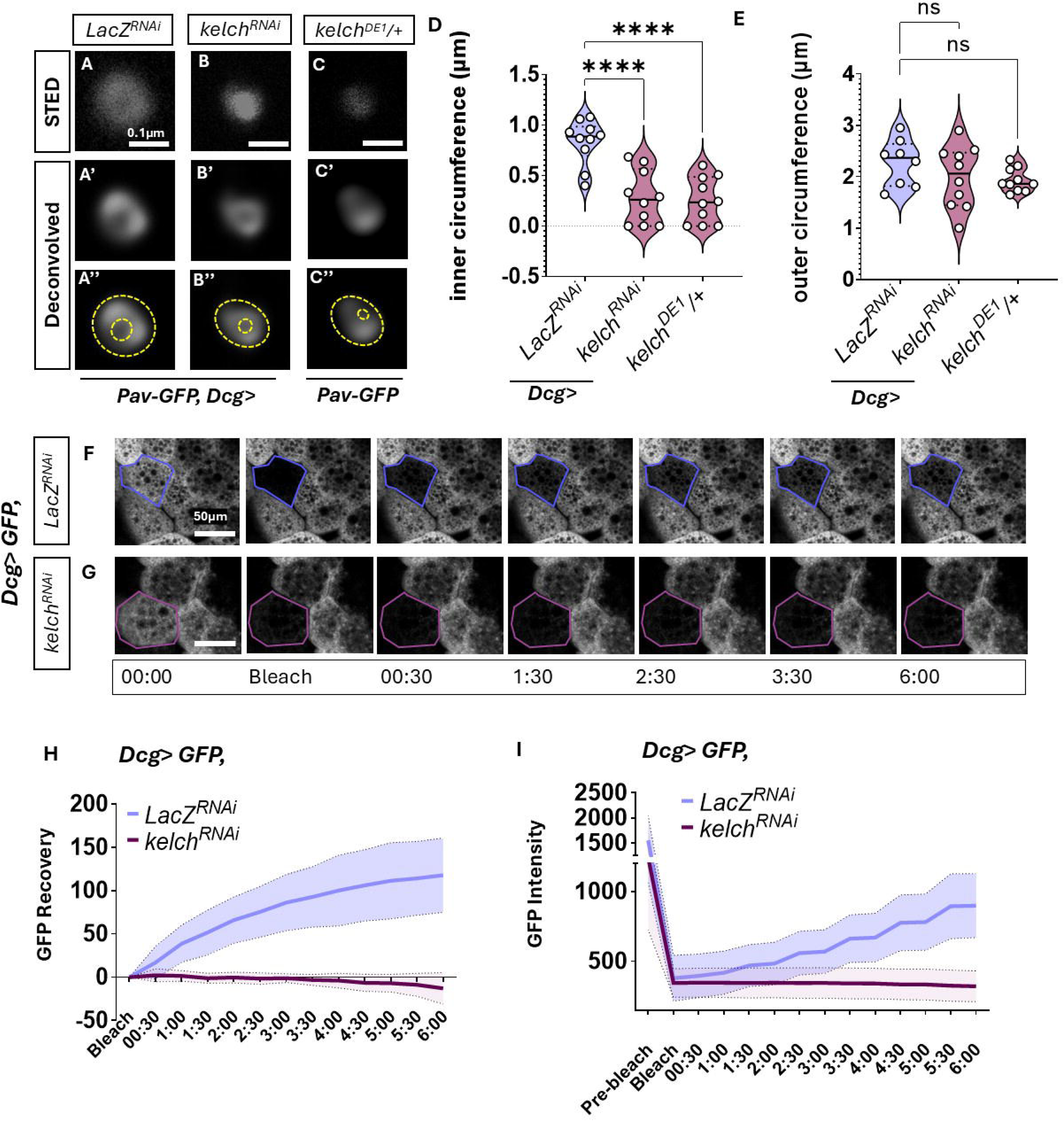
Kelch is required for ring canal structure and function. A-C. Representative high magnification confocal image acquired as in 1G-H of pav-GFP with STED resolution (A-C) and Hugyens deconvolution (A-C’) from 112h AED adipocytes expressing a control *LacZ^RNAi^* (A), *Kelch^RNAi^* (B) or from *Kelch* heterozygotes (*Kelch^DE1^/+,* C). RNAi expression was driven using *Dcg-GAL4.* A”-C” are images from A’-C’ with inner and outer circumference of the ring canal marked in yellow dotted lines. D-E. Measurement of average inner (D) and outer circumference (E) from A-C. n= 8-10 ring canals from at least five different animals per sample F-G. Time lapse montage over six minutes of fluorescent images showing GFP expression driven by *Dcg-GAL4* in FRAP experiments as in 3A-B from 112h AED adipocytes also expressing *LacZ ^RNAi^* (F) or *Kelch ^RNAi^* (G). The bleached cell is outlined in blue in F and magenta in G. H. Quantification of percent GFP recovery in bleached cells from F-G. Data represent average of 10 independent photobleaching experiments from ten animals collected across 3 independent crosses, standard deviation is plotted as the shaded error envelope. Scale bars: A-B = 50µm; F-G = 0.1µm

We next tested whether disruption of ring canal integrity using *Dcg>Kelch^RNAi^*impacted intercellular cargo transport. We performed FRAP experiments as in **Fig. 2A** to observe recovery of cytoplasmic GFP after photobleaching in control and *Dcg>Kelch^RNAi^*animals. We observed no recovery in GFP fluorescence in six minutes in *Dcg>Kelch^RNAi^* adipocytes (**Fig. 3G-I**, whereas control adipocytes showed visible and robust recovery in fluorescence as before (**Fig. 3F, H-I**). These results establish that Kelch is a necessary component of the fat body ring canal, disrupting which severely impairs the rapid transport of cytoplasmic proteins through ring canals.

### Ring canal disruption results in increased ER stress and uncoupling of adipocytes

With a suitable method to disrupt ring canals in the fat body using *Dcg>Kelch^RNAi^*, we sought to test our hypothesis that disrupting ring canal structure negatively impacts the ER stress response in the fat body. To interrogate the effects of *Kelch* knockdown on ER stress tolerance, we used the 4EBP^intron^-DsRed reporter [Kang et al., 2017]. We found that knocking down *Kelch* in 112h AED adipocytes resulted in higher levels of DsRed expression, suggesting that these animals are experiencing higher levels of stress in adipose tissue than control *Dcg>LacZ^RNAi^* animals (**Fig. 4A-B, D**). Notably, these fat bodies do not show disparities in the expression of a control *UAS-GFP* also driven by *Dcg-GAL4* (**Fig. 4A”-C”, E**), indicating that the differences in DsRed reporter levels are not due to global protein synthesis defects.

**Figure 4:**
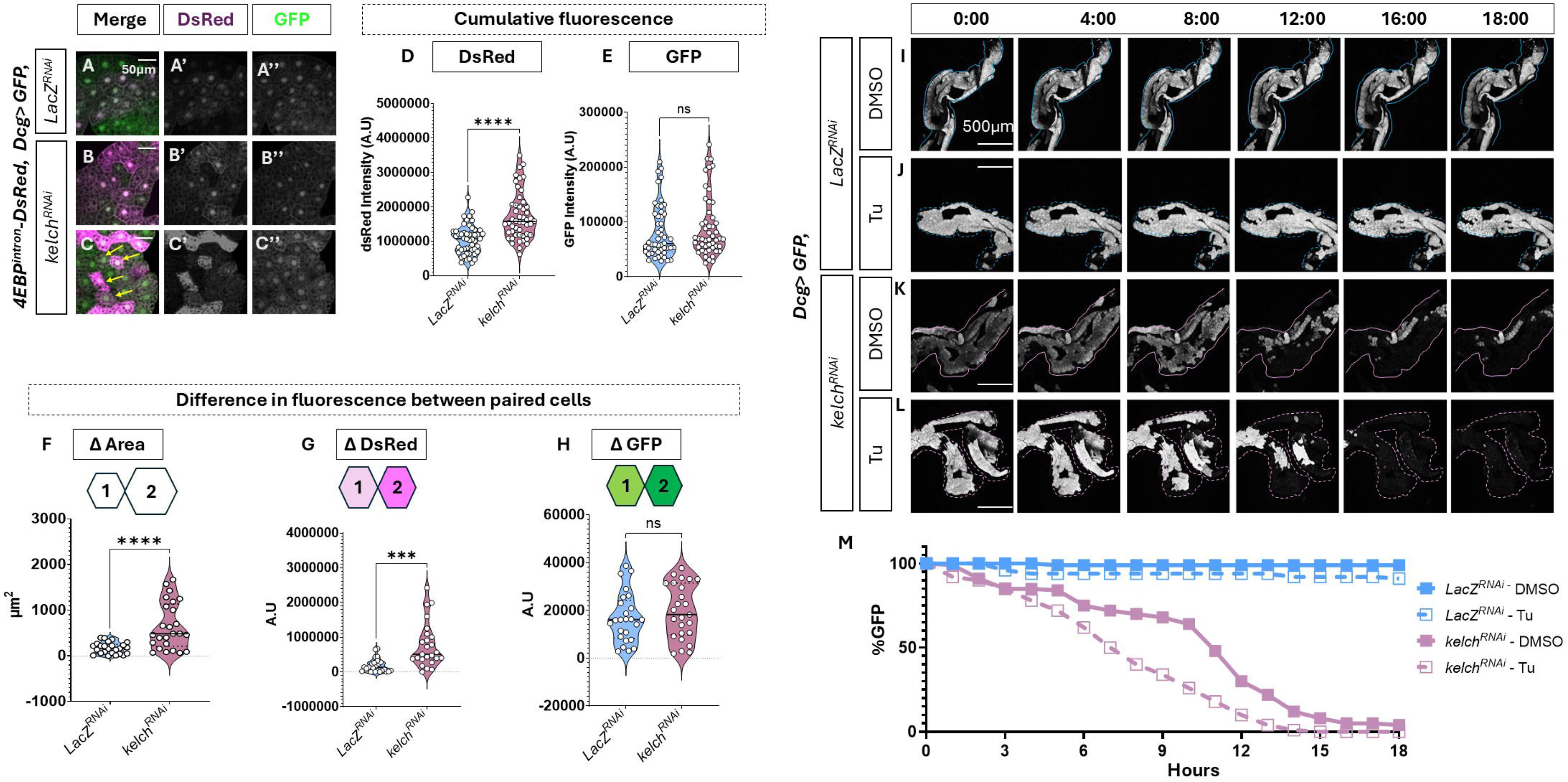
Disruption of ring canals increases susceptibility to ER stress. A-C. Representative confocal images of 112h AED adipocytes expressing 4EBP^intron^-DsRed (Magenta) and GFP (green) driven by *Dcg-GAL4*, which is also driving expression of control *LacZ^RNAi^* (A) or *kelch^RNAi^*(B-C). C represents example of very high DsRed expression in *kelch^RNAi^* (see text). D-E. Quantification of DsRed (D) and GFP (E) signal from A-C. n=50 cells for each genotype, from images taken on fat bodies from at least five different animals across 2 independent crosses. F-H. Measurement from ring-canal paired 112h AED adipocytes cells in A-C showing differences in absolute values of 4EBP^intron^-dsRed intensity (F), cell area (G) and GFP intensity (H). n=25 pairs for each genotype from five animals across 2 independent crosses. I-L. Time lapse fluorescence image montages over 18 hours of GFP expression (green) in 112h AED fat bodies treated with either vehicle (I, J) or Tunicamycin (J, L). *Dcg-GAL4* drives GFP expression and expression of *Lac^RNAi^*(I-J) or *Kelch ^RNAi^* (K-L). Please note that these images are at lower magnification showing a larger section of the fat body. M. Quantification of GFP signal from I-L. Scale bars: A-C = 50µm; I-L = 500µm

Interestingly, in some *Dcg>Kelch^RNAi^* fat bodies, we also observed some cells displaying highly disparate DsRed expression between paired adipocytes (**Fig. 4C, C’**), which contrasts with our original observation that putatively paired adipocytes show similar levels of DsRed reporter under normal conditions (**Fig. S1, 4A, A’**). This prompted us to ask whether ring canals might mitigate or “buffer” ER stress across adipocytes. To understand this, we performed pairwise measurements in all fat bodies from **Fig. 4A-C** to compare differences in control GFP, 4EBP^intron^-DsRed, and cell size, between paired adipocytes. We found that while the difference GFP intensity did not vary between control and *Kelch^RNAi^* expressing adipocytes (**Fig. 4F)**, DsRed fluorescence, as well as the difference in cell area between paired cells did show a significant difference (**Fig. 4G-H**). These results collectively suggest a critical role for ring canals in maintaining homeostatic tissue function by equilibrating protein levels and thus buffering ER stress levels across paired cells.

### Disrupting ring canal structure results in lowered survival in stressed fat body

Since we found that *Kelch^RNAi^* expressing fat bodies display higher levels of ER stress, we next investigated a role for ring canals in the response to acute stress. To assess the stress-responsiveness of fat body to extrinsic stressors in real time, we developed an *ex vivo* live imaging protocol that allows for long-term monitoring of adipose tissue in the presence of ER stress-inducing drugs such as Tunicamycin. This strategy allowed us to monitor the effects of tissue-autonomous effects of exogenous ER stress on fat body. We embedded third instar larval fat bodies from *Dcg>GFP* animals expressing either *LacZ^RNAi^* or *Kelch^RNAi^*in low-melting agarose and incubated them in complete Schneider’s insect medium containing either vehicle (Veh = DMSO) or Tunicamycin. This experimental set up allowed for time-lapse imaging over 18 hours to assess adipocyte viability by measuring the irreversible loss of GFP. While *Dcg>LacZ^RNAi^*fat bodies treated with DMSO remained viable for the entire time course (**Fig. 4I, M**), comparably treated *Dcg>Kelch*^RNAi^ fat bodies rapidly lost GFP expression in cells rapidly, with up to 50% of cells losing GFP expression within 11 hours, and nearly 95% in 18 hours (**Fig. 4K, M**). This loss of viability was even more pronounced in *Kelch^RNAi^*fat bodies that were treated with Tunicamycin, where over 50% of cells lost GFP expression within six hours and complete GFP loss at 13 hours (**Fig. 5L, M**). In comparison, the *Dcg>LacZ^RNAi^* fat bodies showed a gradual loss in viability, with only 2% cells losing GFP expression over the time course of 18 hours (**Fig. 5J, M**). Taken together, these findings corroborate our hypothesis that ring canal-mediated transport results is required for ER stress buffering in fat bodies, and mediates cell survival under conditions of very high extrinsic stress.

**Figure 5:**
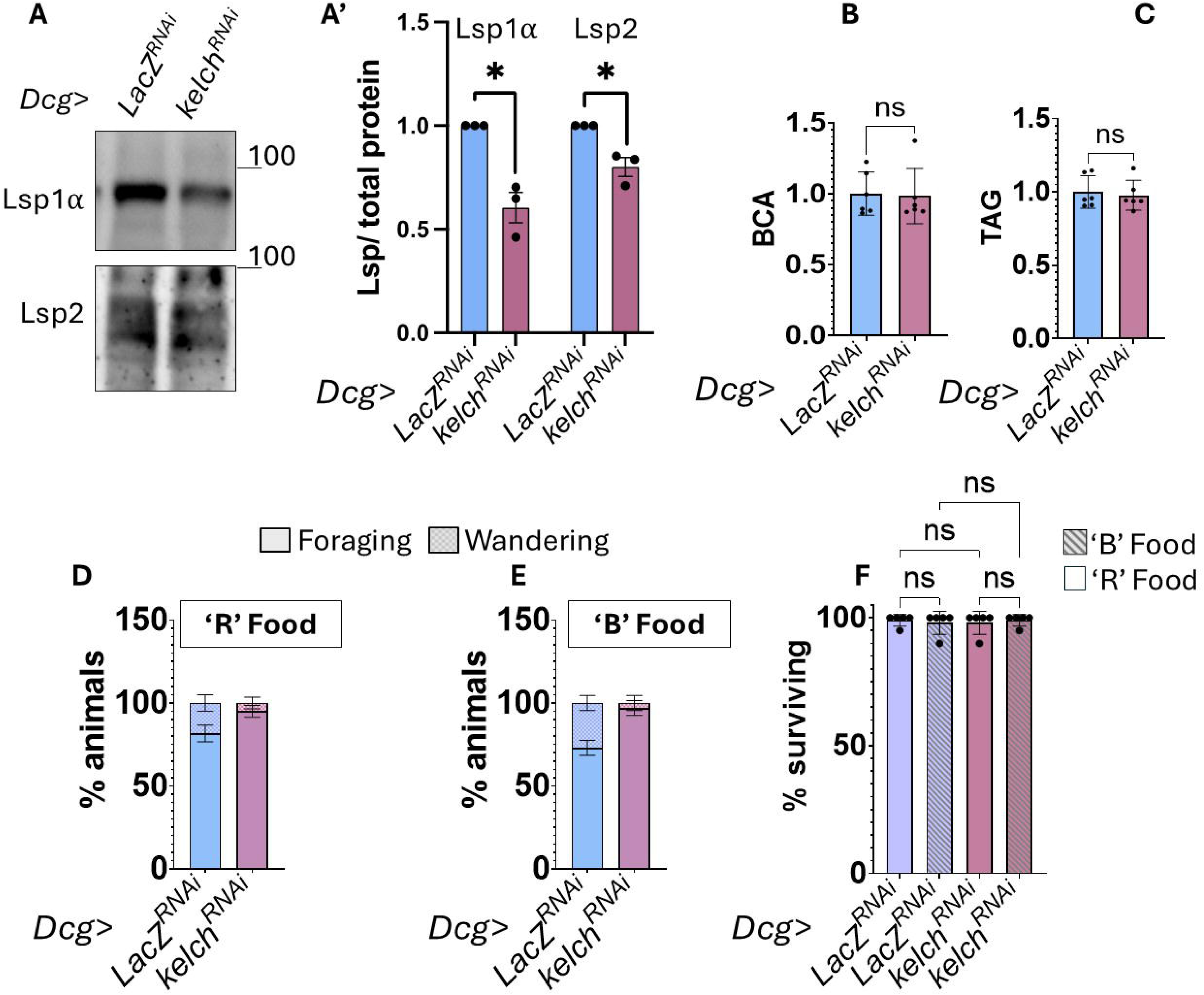
Disruption of ring canals reduces protein secretion from the fat body and results in delayed development. A. Western blot analysis of Lsp1α and Lsp2 protein levels in extracts from whole 112h AED larva where *Dcg-GAL4* drives either control *Lac^RNAi^* or *Kelch ^RNAi^*. Data are quantified in A’, bars represent average of 3 independent experiments from two independent crosses, and error bars represent standard error. B. Relative total protein levels measured by BCA assay levels in whole 112h AED larvae where *Dcg-GAL4* drives expression of either *LacZ^RNAi^* or *Kelch ^RNAi^*. Graph represents the average of six biological replicates and error bars indicate standard deviation. C. Relative triacylglyceride (TAG) levels in animals from B. D-E. Developmental timing as measured by the percentage of larva either foraging or wandering at 112h AED when raised on protein rich food (‘R’) or basic food (‘B’). *Dcg-GAL4* drives either control *Lac^RNAi^* or *Kelch ^RNAi^*. n=5 vials per genotype with 25 larvae per vial (see methods). F. Percentage of animals that survive to adulthood from D-E.

### Disrupting fat body ring canals impacts secretory capacity of adipocytes and larval developmental timing

Given that the fat body is a critical tissue for proper development, and that it relies on ER stress response pathways for its homeostatic function [Huang et al., 2017; Kang et al., 2017; Sone et al., 2013]; we sought to determine the phenotypic consequences of disrupting ring canals in adipocytes. The primary function of the fat body particularly during the third instar stage is to assimilate dietary nutrients from the hemolymph, which is then used to 1) to synthesize, store, and secrete larval serum proteins (Lsps) back into the hemolymph [Powell et al., 1984; Roberts et al., 1977], and 2) synthesize and store lipid reserves in the form of triacylglycerides in adipocytes [Grönke et al., 2005]. These proteins and lipids are sequestered in the fat body and hemolymph until later non-feeding stages of development, when they are broken down to support the energetic needs of metamorphosis [Heier and Kühnlein, 2018; Jowett et al., 1986] . We asked whether the high levels of ER stress, as well as disparate ER stress reporter between sister cells seen with ring canal disruptions (**Fig. 4A-H**) impacted protein secretory capacity or lipid storage in *Dcg>Kelch^RNAi^* animals. We first examined protein secretory capacity by western blotting analysis of two larval serum proteins, Lsp1α and Lsp2, in whole larvae extracts from either *Dcg>LacZ^RNAi^* or *Dcg>Kelch^RNAi^* animals. We found that the levels of both Lsp1α and Lsp2 were significantly lower in *Dcg*>*Kelch^RNAi^*(**Fig. 5A-A’**) animals, suggesting that disrupting ring canals likely impairs the secretory function of adipocytes, which could impact development. However, total protein levels were unchanged in *Dcg>LacZ^RNAi^* and *Dcg>Kelch^RNAi^* animals (**Fig. 5B**) . We next tested whether disrupting ring canals in the fat body resulted in lipid storage defects. The majority of lipids are stored in the form of triglycerides in *Drosophila.* To measure potential defects in lipid storage, we performed triglyceride assays on whole larval extracts from

*Dcg>LacZ^RNAi^* or *Dcg>Kelch^RNAi^*animals. Interestingly, we observed no difference in the amount of total triglyceride content between the two genotypes (**Fig. 5C**), suggesting that the ER stress defects observed in *Dcg>Kelch^RNAi^*(**Fig. 4A-H**) impacts fat body protein secretory capacity but not lipid metabolism.

As described above, larval serum protein secretion from the fat body is critical for timely metamorphosis and since *Dcg>Kelch^RNAi^* showed defective Lsp secretion, we examined the developmental timing of these animals. Control larvae reared on a protein-rich diet (‘R’ food) begin wandering around ∼110h AED as expected, but *Dcg>Kelch^RNAi^* larvae displayed a modest but significant delay in developmental timing when reared on a regular diet (‘R’ food, **Fig. 5D**) as evidenced by a much lower population of animals wandering at the same time point. Next, since developmental timing is sensitive to protein availability and we found that disrupting fat body ring canals resulted in reduced protein secretion (**Fig. 5A-A’**), we performed these experiments in animals reared on a reduced protein food (‘B’ food, see Methods). Interestingly, the developmental delay in *Dcg>Kelch^RNAi^* animals was even more pronounced on ‘B’ food (**Fig. 5E**), suggesting an additional role for ring canals in buffering nutrient deprivation stress in addition to ER stress. Notably, the disruption of ring canal structure did not result in any larval lethality (**Fig. 5F**), indicating that ring canals might contribute to the general robustness of larvae, but are not necessary for their survival

Together, these phenotypic analyses underscore the physiological importance of ER stress buffering across neighboring cells by ring canals in fat body function.

## Discussion

In this study we describe the first instance of intercellular bridges in adipose tissue in any organism and establish a function for them in this tissue. We demonstrate a role for specialized sharing of cytoplasmic and secretory pathway contents across adipocytes paired by ring canals (also known as intercellular bridges in vertebrates) in the metabolic and secretory larval fat body, a tissue with several critical functions. Our findings point to a role for rapid intercellular transport of specific organellar cargo in buffering the high levels of intrinsic ER stress adipocytes endure due to their secretory load, and also mediating adipocyte survival under conditions of extrinsic stress. Further, we find that the proper structure and function of ring canals are required for the normal secretory function of the fat body, and that their disruption results in developmental defects. This underscores the importance of intercellular communication in normal adipocyte function. Integrating these observations, we postulate that buffering ER stress across cells is necessary in secretory and metabolic cell types for both normal tissue function and survival under acute stress The broader implications of our findings are discussed below.

### The nanoscale dimensions of ring canals confer functional and cargo specificity

Our super-resolution imaging shows that ring canals in the *Drosophila* larval adipocytes are extremely small but open structures, having an inner diameter of less than 500 nm even at developmental stages when adipocytes span several tens of microns (**Fig. 1**). This means that adipose ring canals are up to two orders of magnitude smaller than the cell. Our work also shows that the ring canals adipose do not grow even as adipocyte nuclei undergo rounds of endoreplication to reach a final average ploidy of ∼255[Nordman et al., 2011]. This is in contrast to other ring canal structures, for example ring canals of nurse cells in the female germline grow in scale with the nuclei [Cooley, 1998]. Although both adipocytes and nurse cells are highly secretory, this contrast in their respective ring canal sizes can be partly explained by the differences in tissue architecture and the role ring canals play in these respective cell types. In the germline, ring canals permit the transport of mRNAs and proteins synthesized in the nurse cells to be deposited into the developing oocyte in an efficient manner [Bernard et al., 2018; Hsu et al., 2015; McLean and Cooley, 2014; Petrella et al., 2007; Roth and Lynch, 2009; Shaikh et al., 2024]. The structure of the ovarian follicle is such that not every nurse cell makes contact with the oocyte, and thus ring canals are the primary mode of intercellular transport from distal nurse cells to the oocyte. However, the larval fat body is a relatively flat monolayer of adipocytes that secretes directly into the circulating hemolymph that makes contact with the entire tissue. Adipocytes hence rely on canonical vesicular trafficking via the secretory pathway to deposit proteins and lipids into the hemolymph[Ugrankar-Banerjee et al., 2023; Yang et al., 2021]. Thus, unlike in the nurse cells, fat body ring canals appear to be cargo selective, likely restrained by their size- this is supported by our finding that they only permit the transport of some cytoplasmic, ER and Golgi proteins, but not mitochondria (**Fig. 2**). However, despite differences in their functions, ring canals in both nurse cells and adipocytes appear to share core components for maintaining their structural integrity such as Kelch (**Fig. 3**). The structural components of ring canals in adipocytes likely will inform their cargo selectivity, in addition to specifying their size. Thus, future studies on larval adipocyte ring canal composition and comparative analyses with other ring canal structures may reveal the mechanistic basis for tissue-specific cargo selectivity.

### A potential fusome-like structure in the larval fat body

Similar to how the cytoplasm is continuous across adipocytes paired by open ring canals (**Fig. 2A-B**), it may be the ER is also contiguous across paired cells, forming potential fusome-like structures that would allow for efficient protein sharing between cells. The rapid recovery (< two minutes) in KDEL-GFP after photobleaching in our FRAP paradigm supports such a speculation (**Fig. 2C**). Since adipocytes are reliant on their ER network for many functions [Yang et al., 2021], it is possible that ring canals may also play a role in regulating ER architecture and vice versa. Recent work in the *Arabidopsis thaliana* root endodermis has revealed that ER structure during cell division influences the formation of plasmodesmata (as intercellular bridges are known in plants), and that ER networks span multiple cells connected through plasmodesmata [Li et al., 2024]. Given the nutrient uptake function of the root endodermis cells, a function also performed by the *Drosophila* fat body, it is tempting to extrapolate that such ER networks could also span across the adipocytes through ring canals, and membrane tethered ER may even regulate the formation of ring canals, as in *Arabidopsis*. Future studies investigating how ER morphology changes during cell division and incomplete cytokinesis could address the potential relationship between this ER structure and ring canal formation in metazoans.

### Adipocytes buffer ER stress through ring canals

The fat body functions to secrete several important proteins into the hemolymph that are required for supplying nutrients to all the developing tissues of *Drosophila* in larvae[Musselman and Kühnlein, 2018]. The fat body also simultaneously functions in the absorption and storage of nutrients as lipids which are required for the animal to successfully undergo metamorphosis. The ER is both the primary site of lipid homeostasis and a crucial part of the protein secretory pathway, and consistently we and other have demonstrated that pathways required for maintaining ER homeostasis are constitutively active in the larval fat body [Huang et al., 2017; Kang et al., 2017; Sone et al., 2013]. The importance of this organelle in fat body function is further emphasized by phenotypes seen in ER stress response pathway mutants in vertebrates and *Drosophila* metabolic tissues [Ho et al., 2018; Moncan et al., 2021]. We find that compromising ring canals by knocking down *Kelch* in the fat body results in higher levels of average ER stress reporter expression, indicative of general ER malfunction (**Fig. 4A-D**). Consistent with such a malfunction, we observe reduced secretion of Lsps in these animals (**Fig. 5A**) which likely form the basis of the nutrient-dependent developmental delay we observed (**Fig. 5D-E**). Notably, since loss of Lsp2 alone has been demonstrated to accelerate development [Valzania et al., 2024], we surmise that disruption of ring canals impact the secretion of many proteins in addition to Lsp1 and 2. More intriguingly, we also find the disrupting ring canals led to the “unpairing” of adipocytes such that sister cells now showed significant differences in cell size and ER stress levels (**Fig. 4F-H**). These results lead us to consider that one of the critical functions of ring canals is to mitigate high levels of ER stress in individual cells by facilitating rapid sharing of cytoplasmic/ER/Golgi contents. This is supported by other work demonstrating that ring canals in somatic ovarian follicle cells serve to equilibrate protein levels across cells connected through ring canals [Airoldi et al., 2011; McLean and Cooley, 2013], though the needs for such equilibration was not fully understood. We propose that ring canal-mediated equilibration in secretory tissues such as the fat body and follicular epithelium serves to coordinate ER stress levels between connected cells such that no single cell experiences the detrimental effects of chronic ER stress response activation. Our data showing that disrupting ring canals in fat bodies subjected to exogenous stress results in rapid adipocyte death (**Fig. 4I-M**) strongly supports the hypothesis that ring canals serve to buffer ER stress in sister cells. An open question for future studies is whether such stress buffering is mediated by transfer of specific proteins or molecules. Further, our study has only tested the role ring canals may play in buffering ER stress. It would be interesting to investigate whether and how ring canals play a role in modulating or buffering other stresses such as mitochondrial, lipotoxic, osmotic or oxidative stressors. Finally, recent work from our lab shows a role for stress response in sexual dimorphic stress tolerance in larvae[Grmai et al., 2024]. While all the experiments performed in this study are in female larvae, it would be interesting to determine whether the function of ring canals may contribute differentially to larval adipose sexual dimorphism in future studies[Grmai et al., 2024].

### Ring canals allow adipocytes to selectively couple secretory function

As described above, the larval fat body has both lipid metabolism and protein secretion activity, which relies on proper ER function. Dietary lipids and nutrients absorbed the by the fat body are stored as triacyl glyceride incorporated into lipid droplets which are synthesized on the surface of the ER [Wilfling et al., 2014]. Our results show that disruption of ring canals leads to reduced total serum protein levels (**Fig. 5A**) but does not impact the levels of triglycerides stored in larvae (**Fig. 5C**), indicating that ER stress equilibration via fat body ring canals may impact protein secretory capacity, but not lipid metabolism. The majority of lipids in the larval fat body is stored in the form of lipid droplets and previously published literature suggests that even the smallest lipid droplets (∼10µm) might not be large enough to pass through ring canal openings since they are larger than the opening of ring canals (1-2µm) [Fan et al., 2017]. Together, these analyses suggest that ring canals facilitate the selective coupling of adipocyte protein secretory functions, without impacting lipid storage functions. Thus, ring canals could lend the fat body an elegant mechanism to separate nutrient dissemination functions, which occur rapidly and constantly throughout development, from equally important anabolic lipid storage functions, which is a more passive storage function critical for nutrients later during metamorphosis. It is worth mentioning here that the larval fat bodies present a range of lipid droplet sizes [Ugrankar et al., 2019], some of which may be well below the detection capabilities of our live imaging experimental set up. Further, it is also possible that lipid metabolic defects in *Kelch* loss of function fat bodies manifest as lower levels of glycogen, another major energy storage moiety in the larval fat body [Garrido et al., 2015].

### Does multinucleation confer advantages to metabolic tissues?

Paired sister cells in the fat body do not undergo complete cytokinesis, and remain connected to one another partially sharing cytoplasm, making each pair a bi-cellular syncytium. Syncytia or syncytial structures form either through cell fusion or through incomplete cell cycles resulting in multiple nuclei sharing cytoplasm[Peterson and Fox, 2021]. It is tempting to propose that the extent of intercellular transport and cargo sharing might even render larval adipocyte pairs binucleate. Interestingly, the adult fat body of *Drosophila* comprises binucleate and tetranucleate cells which form as a result of cell fusion of a different set of progenitor cells during late metamorphosis[Lei et al., 2022]. However, we find that adult adipocytes do not possess ring canals (data now shown). Both the larval and the adult fat body, therefore, contain syncytial cells- the larval fat body comprising several bi-cellular (or binucleate) syncytia, and the adult fat body containing binucleate and tetranucleate syncytial cells[Lei et al., 2022]. Thus, context dependent variation in DNA content and nuclear number appears to be an important facet of *Drosophila* adipocyte physiology. Multinucleation has also been reported in cultured vertebrate adipocytes [Xu et al., 2017], and in several metabolically active tissues such as cardiomyocytes[Gan et al., 2020], liver hepatocytes[Fortier et al., 2017], mammary epithelial cells[Rios et al., 2016], and exocrine pancreas[Oates and Morgan, 1989]. Our work adds to the substantial and growing body of literature demonstrating the diversity of variant cell cycle processes resulting in tissues containing cells with polyploid DNA content, and binucleate cells [Nandakumar et al., 2021]. However, despite the descriptions of multinucleation in various tissues across phyla, our understanding of the functions and consequences of having multiple nuclei in these vastly different tissues remains largely speculative: However, emerging work present tantalizing hints that multinucleate cells, specifically cardiomyocytes and enterocytes, show improved cellular function in comparison to their mononucleate counterparts [Lam et al., 2025; Lessenger et al., 2025], . Our findings described in this paper thus present *Drosophila* adipose an excellent genetically pliable model to investigate the both the extent and mode of multinucleation in the function of metabolic tissues.

## Materials and Methods

### Fly Rearing

All flies were raised at 25°C. Crosses were set up with ten female virgins and five males no more than three days post-eclosion at the time of setting up the cross. Unless otherwise mentioned, all crosses and rearing were performed on ‘R’ food formulation from Lab Express (https://www.lab-express.com/DIS58.pdf). Crosses were flipped every day to ensure no overcrowding of vials occurred. All fly stocks used in the study are listed in **Table 1**. Specific genotypes for each experiment are listed in **Table 2**.

**Table 1.** List of fly stocks and corresponding stock numbers used in this study.

| Transgene | Source (RRIDs from BDSC) |
| --- | --- |
| *pav-GFP* | BDSC_81650 |
| *UAS-GFP* | BDSC_1522 |
| *UAS-KDEL-GFP* | BDSC_9898 |
| *UAS-Golgi-GFP* | BDSC_30902 |
| *UAS-mito-GFP* | BDSC_8442 |
| *Dcg-GAL4* | BDSC_7011 |
| *UAS-LacZ<sup>RNAi</sup>* | [Kang et al., 2017] |
| *UAS-Kelch<sup>RNAi</sup>* | BDSC_31251 |
| *Kelch<sup>DEI</sup>* | BDSC_4893 |
| *UAS-GFP<sup>NLS</sup>* | BDSC_4776 |
| *4EBP<sup>intron</sup>-DsRed* | [Kang et al., 2017] |
| *4EBP<sup>intron</sup>-GFP* | [Grmai et al., 2024] |

**Table 2.** List of genotypes by figure.

| Figure | Genotype |
| --- | --- |
| S1 | *;Dcg-GAL4,UAS-GFP<sup>NLS</sup>/UAS-LacZ<sup>RNAi</sup>;4EBP<sup>intron</sup>-DsRed/+* |
| 1 | *pav-GFP* |
| 2A-B | *;Dcg-GAL4/UAS-GFP;* |
| 2C | *;Dcg-GAL4/UAS-KDEL::GFP;* |
| 2D | *;Dcg-GAL4/+;UAS-Golgi-GFP/+* |
| 2E | *;Dcg-GAL4;UAS-mito::GFP;* |
| 3A | *;Dcg-GAL4,pav-GFP/UAS-LacZ<sup>RNAi</sup>;* |
| 3B | *;Dcg-GAL4,pav-GFP/+;UAS-Kelch<sup>RNAi</sup>/+* |
| 3C | *;Kelch<sup>DEI</sup>/pav-GFP;* |
| 3F | *;Dcg-GAL4/UAS-LacZ<sup>RNAi</sup>;* |
| 3G | *;Dcg-GAL4/+;UAS-Kelch<sup>RNAi</sup>/+* |
| 4A, I-J | *;Dcg-GAL4,UAS-GFP<sup>NLS</sup>/UAS-LacZ<sup>RNAi</sup>;4EBP<sup>intron</sup>-DsRed/+* |
| 4B-C, K-L | *;Dcg-GAL4,UAS-GFP<sup>NLS</sup>/+;4EBP<sup>intron</sup>-DsRed/ UAS-Kelch<sup>RNAi</sup>* |
| 5A-F | *;Dcg-GAL4,UAS-GFP<sup>NLS</sup>/UAS-LacZ<sup>RNAi</sup>;4EBP<sup>intron</sup>-DsRed/+*<br>*;Dcg-GAL4,UAS-GFP<sup>NLS</sup>/+;4EBP<sup>intron</sup>-DsRed/ UAS-Kelch<sup>RNAi</sup>* |

### Staining for confocal microscopy

Larval fat bodies were micro dissected in 1X phosphate buffer saline (PBS) and fixed in 4% paraformaldehyde (PFA) made fresh to a final concentration of 1X PBS for 25 minutes on a nutator. Tissues were washed three times each in 0.1% PBS-Tween between different stains. 4′,6-diamidino-2-phenylindole (DAPI, 200 μM final) and optional phalloidin Alexafluor 647 (Thermo Fisher Scientific, 1:1000) staining was performed to counterstain for nuclei and actin/cell membrane respectively for 1h at room temperature. Tissues were mounted on glass slides in a solution of 70% glycerol prior to placing coverslips. Slides were sealed with a thin coating of nail polish to prevent evaporation. If protected from light, slides are stable at room temperature for a month, at 4°C for several months and at -20°C for several years with limited reduction in fluorescence of GFP, RFP and common fluorescent dyes.

### Super resolution microscopy

Fat bodies mounted in 70% glycerol without DAPI were imaged using a 93x glycerin immersion objective (refractive index-matched) on a Leica-SP8 imaging system. Each ring canal image was captured as a z-stack with 10 steps (0.1-0.2µm step size, system optimized conditions used) 3D STED imaging of pav-GFP marked ring canals was performed using a 592mm wavelength depletion laser. Further deconvolution was performed using the Huygens suite with the same iteration settings across all experimental conditions. Super-resolved images were then used to measure ring canal dimensions by manually tracing inner and outer circumferences using ImageJ.

### Fat body mounting for live imaging

Fat bodies dissected in 1X PBS were carefully mounted on MatTek glass bottom culture dishes in a solution of 1% low melting agar in 1X PBS as follows. A 3X agar stock solution was melted at 90°C and diluted with chilled PBS to bring the working agar solution to 25°C. 250µl of agar was carefully pipetted on the glass bottom dish divot in which fat bodies are placed within two minutes. After agar gelling occurs (5 minutes), specimens are immediately imaged on a Nikon A1 inverted line scanning confocal microscope.

### Fluorescence recovery after photobleaching (FRAP) experiments

FRAP imaging was performed using a 20X dry objective on a Standard Nikon A1 plus Confocal with the pinhole opened to allow for increased fluorescence detection of 3D mounted tissues at a depth of up to 25µm. Individual cells were traced using the polygon ROI function. Standard Nikon A1 plus stimulation settings were used to deplete fluorescence reporters (100% laser power for depletion, 1/12 scanning speed, four rounds of stimulation to achieve 95% or higher depletion in total fluorescence of ROI of interest). Processing and quantifications of all FRAP experiment images were performed using the NIS-elements advanced microscopy suite. A custom GA3 workflow pipeline was developed to perform timed image quantifications, which were exported to excel spreadsheets and graphed using Graphpad prism.

### SDS-PAGE and Western Blotting

Whole larvae (3 per sample) were flash frozen before being crushed using a pestle in 125µL of Radio-immunoprecipitation assay lysis buffer (RIPA, 50 mM Tris HCl, pH 7.4, 150 mM NaCl, 1% Triton X-100, 0.5% Sodium deoxylcholate, 0.1% SDS, 1 mM EDTA) supplemented with protease inhibitor (Roche). The lysate was centrifuged at 16000 RCF for 10 minutes, and clear supernatant was transferred to a new tube while avoiding the floating fat layer. Total protein was measured using 5μl lysate in a standard BCA assay in a plate reader. 25μl of lysate mixed with 4X Laemmli buffer containing 4% β-mercaptoethanol was analyzed by SDS-page followed by western blotting onto nitrocellulose membranes. Membranes were blocked with 5% milk in 1xPBS containing 0.1% Tween-20 and then probed overnight at 4°C with Rat anti-Lsp1α (1:1000) and Lsp2 (1:2500) [Valzania et al., 2024]. Proteins were detected using Rat anti-HRP secondary antibody. Blots were quantified in ImageJ by measuring intensity of specific bands and normalizing it to BCA measurements.

### Measurement of developmental delay

Crosses with 20 virgins and 10 males were set up in population cages with a plate of grape agar supplemented with fresh yeast paste. Crosses were flipped after 1h egg lays. 24 hours after egg lays were complete, 25 1st instar larvae of similar size were picked from grape plates and placed in vials containing either ‘R’ (regular/protein rich) or ‘B’ (Bloomington formulation, https://bdsc.indiana.edu/information/recipes/bloomfood.html) food, which contain 4% yeast w/v and 1.5% yeast w/v respectively. To determine percentage of wandering larvae, measurements were made at 112h AED (after egg deposition). Total survival to adulthood was measured by counting animals post eclosion from individual vials.

### TAG assay

3 whole larvae per sample were crushed in a solution of 0.5%SDS-1X PBS, incubated on ice for 15 minutes, and then centrifuged at 12000 RCF at 4°C for five minutes. 3µL of the lysate was added to 100µL of Triglyceride reagent solution in 96 well dish wells, in technical duplicates. After 15 minutes of incubation, the plate was analyzed on an absorbance plate reader at 540nm. Total protein was simultaneously measured using 5μl lysate in a standard BCA assay in a plate reader.

### Statistical analyses

The figure legends describe the sample sizes for the corresponding data. Significance in all graphs were calculated using unpaired T-test with Welch’s correction for unequal standard deviations.

## Acknowledgements

We would like to thank publicly available model organism resources that fueled our research: Flybase, Drosophila Ortholog Prediction tool (DIOPT) and Bloomington *Drosophila* stock center. We would like to thank all members of our lab for discussion and feedback on the project. We also extend our thanks to Drs.Lydia Grmai, Yang Hong, and Lesley Weaver for critical discussions and feedback on the manuscript. Anti-Lsp1 and Lsp2 antibodies were a generous gift from the lab of Dr. Pierre Leopold. Stocks obtained from the Bloomington Drosophila Stock Center (NIH P40OD018537) were used in this study. We used FlyBase (release FB2024_01) for identifying phenotypes and stocks in this study.

---

**Supplemental Figure 1: Heterogeneity in stress reporter expression**

Representative image showing heterogeneity in 4EBP^intron^-DsRed expression occurring in pairs of neighboring cells (yellow arrows).

Scale bar = 100µm

