## Supplementary figures and images for "Ring canals in the larval adipose of *Drosophila* buffer stress response"

### Supp.Fig.1

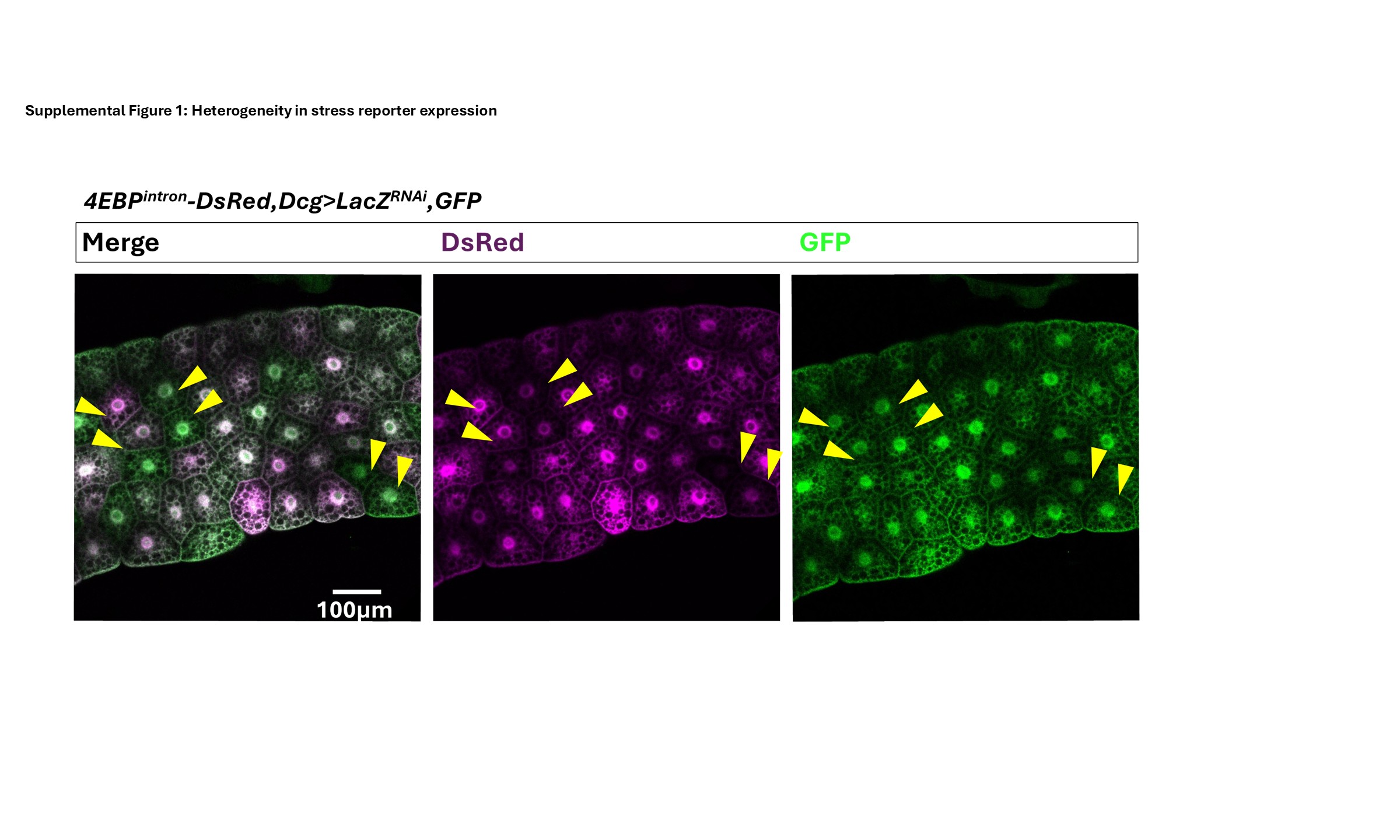
